# Knock-down of genes essential for homoacetogenic growth using sugar inducible promoters in the thermophile *Thermoanaerobacter kivui*

**DOI:** 10.1101/2024.06.18.598388

**Authors:** Benjamin Zeldes, Sabina Mittelstedt, Christoph Baum, Adilia Shakirova, Anja Poehlein, Rolf Daniel, Volker Müller, Mirko Basen

## Abstract

Given recent interest in hydrogen (H_2_) as a medium for storing and transporting renewable energy, the ability of acetogenic bacteria to use H_2_ to directly fix carbon dioxide (CO_2_) shows great promise. Thermophilic bacteria have long been of interest as industrial strains due to their resistance to contamination. *Thermoanaerobacter kivui* is among the most thermophilic acetogens. It can be genetically manipulated, but as yet only in an all-or-nothing manner, either complete gene knockouts or very strong constitutive promoters leading to protein overexpression. The sugar inducible promoters characterized here provide the finer control necessary to close remaining gaps in understanding of *T. kivui*’s energy metabolism and for controlled expression of recombinant genes of industrial relevance.

Acetogenic bacteria are capable of growing on mixtures of the gasses H_2_, CO_2_, and CO using the Wood-Ljungdahl pathway, the most energy efficient CO_2_ fixation pathway. *T. kivui* is a thermophilic (T_opt_ = 66°C) acetogen also capable of growth on sugars including glucose, mannitol, and fructose. RNA-sequencing indicated an operon of phosphotransferase import genes up-regulated on mannitol, and growth on a mixture of glucose and mannitol was diauxic. Given this evidence for sugar responsive transcriptional regulation, the promoter region from the mannitol operon, and a similar operon involved in fructose uptake were used to control expression of a thermostable β-galactosidase reporter gene. Both promoters resulted in higher reporter gene activity when grown on their inducing sugar relative to glucose. The inducible promoters were used to create “knock-down” mutants of the proton-pumping Ech1 hydrogenase. Ech1 is believed to be essential because repeated attempts to knock-out the gene operon have failed. The resulting sugar-inducible Ech1 expression strains grew better on their inducing sugar than on glucose. The mannitol inducible promoter was also used to control expression of an essential gene of the Wood-Ljungdahl pathway, formyl-THF synthetase (*fhs*), involved in consuming the metabolic intermediate formate. When *fhs* repressed by growth on glucose significantly more formate accumulated in the culture medium than during growth on mannitol, clear evidence of a metabolic bottleneck under the knock-down condition. The promoters characterized here provide finer control of recombinant or native gene expression in *T. kivui*, while the thermostable reporter gene allows for rapid characterization of new promoters.

## Introduction

The thermophilic acetogen *Thermoanaerobacter kivui* is one of the most thermophilic acetogens characterized to date (T_opt_ = 66°C), and is of particular biotechnological interest due to its ability to convert syngas (containing H_2_, CO_2_ and CO) to acetate (Weghoff and Müller 2016). Through metabolic engineering, it should be possible to generate more valuable carbon chemicals from these gasses which would otherwise be burned and released as greenhouse gasses (Bengelsdorf and Dürre 2017). However, in addition to autotrophic growth on gasses, *T. kivui* also exhibits growth on a limited set of simple sugars, including glucose, fructose, mannose (Leigh, Mayer, and Wolfe 1981), and the sugar alcohol mannitol (Moon et al. 2019). Heterotrophic growth on these substrates results in very efficient carbon conversion due to use of the Wood-Ljungdahl carbon fixation pathway to re-incorporate CO_2_ released by fermentation. Mixotrophic growth of acetogens on both sugars and gasses has also been proposed as a way to maximize product yields (Fast et al. 2015; Jones et al. 2016).

Acetogens fix CO_2_ via the Wood-Ljungdahl pathway (WLP, or reductive acetyl-CoA pathway), the most energy-efficient carbon fixation pathway known (Bar-Even, Noor, and Milo 2012). In *T. kivui* most genes of the WLP are clustered together in a large operon starting with formyl-tetrahydropholate (THF) synthetase (*fhs*), which activates CO_2_ - derived formate with THF for further reduction by subsequent genes (Hess et al. 2014). Knocking out one of *Acetobacterium woodii*’s two *fhs* genes resulted in a severe growth defect, and during growth on H_2_/CO_2_ the mutant strain accumulated more formate than the wild-type (Moon et al. 2021). In contrast, deletion of the sole *fhs* gene in *T. kivui* is likely to be lethal, since cells would no longer be able to carry out acetogenesis.

During autotrophic growth electrons for carbon fixation via the WLP derive from H_2_. For energy conservation the WLP must be coupled to a transmembrane gradient via either an Rnf or Ech transmembrane complex (Schuchmann and Müller 2014). The energetics of Rnf-containing acetogens have been elucidated, but gaps remain in our understanding of Ech containing acetogens like *T. kivui*, since with current models their metabolism would yield negative ATP (Kremp, Roth, and Müller 2022). *T. kivui* contains two *ech* gene clusters, which are responsible for coupling WLP activity to energy conservation either individually or in combination. The Ech2 complex has been purified and confirmed to function as a proton-translocating Ferredoxin-dependent hydrogenase (Katsyv and Müller 2022). A knockout strain lacking Ech2 grew the same as the wild-type on H_2_/CO_2_ or sugars, but was unable to grow on CO or pyruvate, suggesting it is essential for ferredoxin recycling under growth conditions in which ferredoxin is the sole redox carrier (Baum et al. 2024). Ech1 is hypothesized to form a complex with and accept electrons directly from MetVF of the WLP (Katsyv et al. 2021), which would make it the key coupling point between the WLP and proton-motive-force energy conservation. The *ech1* gene cluster could not be knocked out after numerous attempts, suggesting that it may be essential. We hypothesized that this essentiality could be overcome by replacing the native promoter with an inducible promoter exhibiting low basal expression (low “leakiness”) under non-inducing conditions to create an Ech1 “knock-down” strain. Comparing growth of the resulting strain under inducing vs non-inducing conditions should help to elucidate the role of *ech1* in *T. kivui* metabolism.

There have been efforts to develop inducible promoter systems in several thermophiles. Some of the first developed sugar-inducible promoters for thermophiles were for *Sulfolobus* species (Berkner et al. 2010), while more recent work has established promoters induced by xylose in *Caldicellulosiruptor bescii* (Williams-Rhaesa et al. 2018), cellobiose (Adalsteinsson et al. 2021) or iron starvation (Fujino et al. 2020) in *Thermus thermophilus*, and various sugars in some *Geobacillus* species (Wada and Suzuki 2020). Inducible promoters are established for several mesophilic acetogens (Flaiz et al. 2022), but so far the only promoters confirmed to function in thermophilic acetogens are constitutive (Bourgade, Minton, and Islam 2021; Hocq et al. 2023). Promisingly, the genes involved in metabolism of mannitol and fructose in *T. kivui* have been identified by gene knockouts, confirming that a mannitol-1-P dehydrogenase gene (*mtlD*, TKV_c02860) is essential for growth on mannitol (Moon et al. 2019), and a 1-phosphofructokinase gene (*fruK*, TKV_c23150) is essential for growth on fructose (Basen et al. 2018). Both occur in operons also containing regulatory genes and components of phosphotransferase system (PTS) sugar-specific EII uptake genes. The PTS system in each operon is presumably involved in mannitol and fructose uptake, and the regulator likely controls expression of the entire operon in response to the presence or absence of the respective sugar.

To quickly characterize promoter strength, fluorescent reporter systems such as the ubiquitous GFP are ideal, but GFP and its derivatives still struggle to function under the combined challenges of thermophilic and anaerobic conditions. Systems which do not require molecular oxygen have been developed, including the Y-FAST protein, which binds small externally supplied fluorophores (Plamont et al. 2016). This system has been successfully applied to mesophilic anaerobes, including acetogens (Flaiz et al. 2022). The Y-FAST system has even been applied in *T. kivui*, but required sub-optimal growth temperatures, and resulted in reduced sensitivity that prevented characterization of all but the strongest promoters (Hocq et al. 2023). Since β-galactosidase is native to several extreme thermophiles, thermostable versions are available and have been used as reporter genes for promoter characterization at or above 70°C (Berkner et al. 2010; Fujita et al. 2015).

The goal of this study was to expand the repertoire of promoters available for recombinant gene expression in *T. kivui*, to allow selective induction only during growth with a specific sugar. A global transcriptional analysis comparing growth on glucose and mannitol was carried out with RNA-sequencing, and expression of the mannitol and fructose gene operons during growth on various sugars was monitored by quantitative RT-PCR. Then the respective promoter regions from these operons were tested, first for their ability to control expression of a thermostable β-galactosidase reporter gene, then to control native genes believed to be essential for acetogenic metabolism. When grown under non-inducing conditions (glucose) these strains exhibited either reduced growth rates or evidence of a metabolic bottleneck, confirming the importance of the “knocked-down” genes for *T. kivui* growth. While *T. kivui* shows promise as a biocatalyst capable of converting hydrogen or carbon monoxide gas into value-added products, to truly realize its potential will require greater control over its native metabolism, as well as any heterologously introduced genes – control which the described inducible promoters could provide.

## Results

### RNA sequencing of glucose and mannitol grown cells

Wild-type *T. kivui* cells were grown to late exponential phase in complex medium containing either 25 mM glucose or 25 mM mannitol, and cells harvested for RNA sequencing. Principle component analysis shows good separation of the two sugar conditions on principle component 1, but one mannitol sample is a clear outlier on component 2 (**Fig. S1**). The mannitol samples were harvested at a slightly higher OD (1.3, vs approximately 1.0 for glucose) and it is possible that this sample had depleted specific medium components or begun the transition to early stationary phase. Despite the outlier, the 19 genes up-regulated on mannitol (log2 fold-change > 2, Padj ≤ 0.001) **Table 1** showed clear evidence of a response specific to this sugar. The mannitol gene operon described earlier (TKV_c02830-60) was up-regulated on mannitol roughly 5-fold (log2-fold = 2.2 to 2.6). Other genes up-regulated on mannitol and likely involved in sugar metabolism included a pyrophosphate-specific *pfp* (Tkv_c18810) alternative to the ATP-dependent phosphofructokinase and a transcriptional regulator annotated as xylose repressor *xylR* (Tkv_c10940). In addition gluconeogenic enzymes phosphoenolpyruvate synthase *ppsA* (Tkv_c10530) and fructose 1,6-bisphosphatase (Tkv_c02390) were up-regulated on mannitol, while a glycogen biosynthesis gene cluster Tkv_c11030-60 just barely missed the log2 cutoff for significance, see **Table S1**. The most strongly up-regulated genes on mannitol were ferrous iron transport proteins A and B (TKV_c01950-60).

**Table 1:**
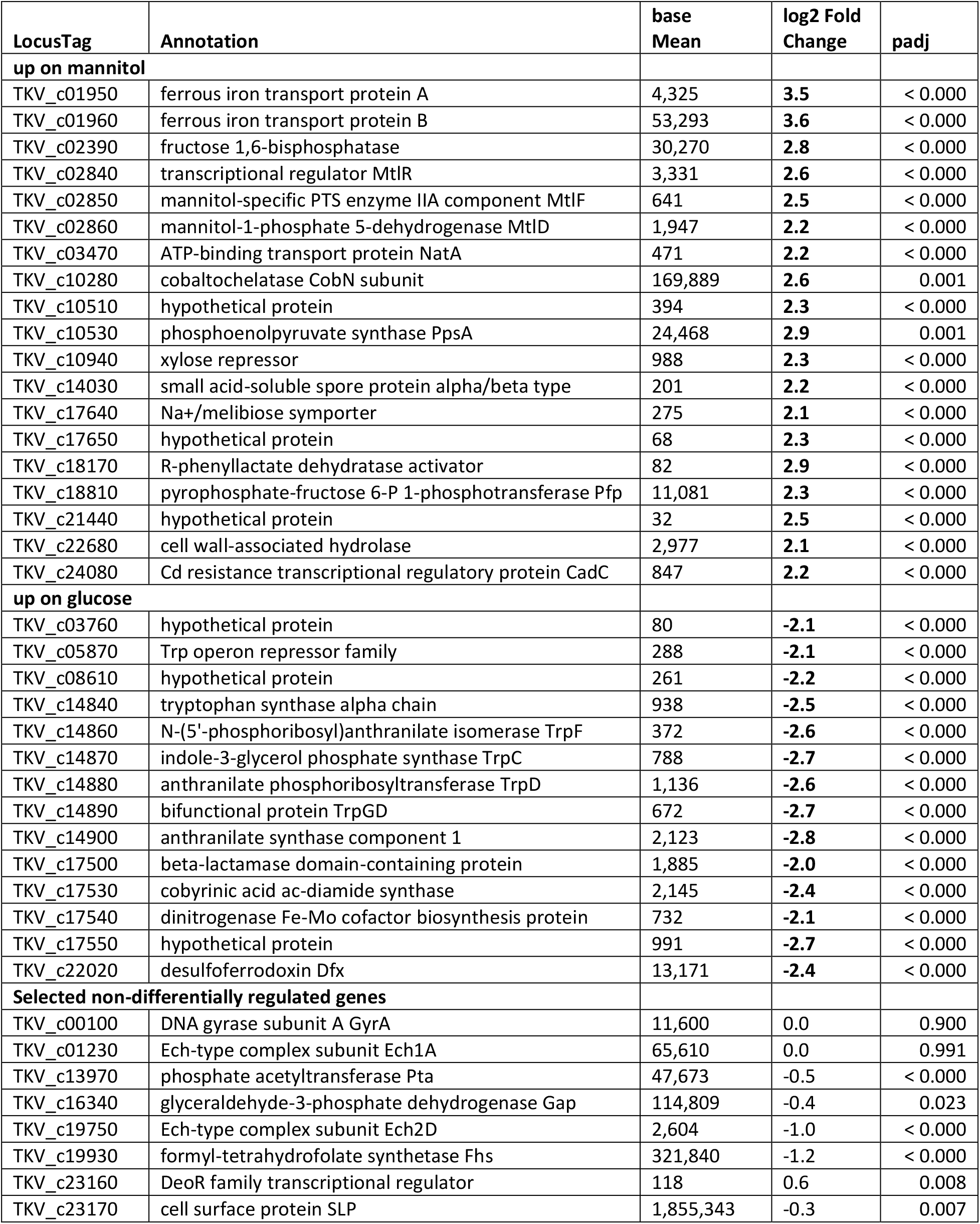
RNA sequencing results comparing growth of wild-type *T. kivui* in complex medium at 66°C on mannitol vs. glucose. Differences of log2 fold-change > 2, with Padj ≤ 0.001 were considered significant.

Among the 14 genes up-regulated on glucose were a cluster of tryptophan biosynthesis genes (Tkv_c14840-90), and parts of a gene cluster annotated as involved in iron-molybdenum cofactor biosynthesis (Tkv_c17530-50). *T. kivui* grows faster on glucose than on mannitol, and the up-regulation of amino acid and cofactor biosynthesis may reflect greater need for these components due to the higher growth rate. Desulfoferrodoxin (Tkv_c22020), which was also up-regulated, can play a role in detoxification of radical oxygen species (Riebe, Fischer, and Bahl 2007), which are generated at higher rates during more rapid growth.

While less quantitative than differential gene regulation, some conclusions can be drawn by comparing relative expression levels (baseMean value in **Table 1**) across genes. Of note here is that, while not differentially regulated, the gene for the S-layer protein (*slp*) (TKV_c23170) is the single most strongly expressed transcript under both conditions, confirming the status of P_slp_ as a very strong constitutive promoter. Transcript levels for the phosphate acetyltransferase *pts* (TKV_c13970), recently described as an even stronger constitutive promoter, were an order of magnitude lower (**Table 1**), while the glyceraldehyde-3 phosphate dehydrogenase *gap* (TKV_c16340) gene fell between these two, although no activity could be detected when using the promoter region for reporter gene expression (Hocq et al. 2023). Key genes of acetogenic metabolism were not differentially regulated but had high baseMean levels, including *fhs* (TKV_c19930), the first gene of the WLP operon, which had the third highest transcript level in the entire genome, and *ech1A* (TKV_c01230), which had slightly more transcript than *pts* according to this data. The genes of the Ech2 operon, starting with *ech2D* (TKV_c19750), were expressed at substantially lower levels than Ech1, as has been reported previously (Baum et al. 2024).

### Growth of *T. kivui* on mixed sugars

*Thermoanaerobacter kivui* grows about equally well on glucose and fructose, but growth is even more robust on mannose (Leigh, Mayer, and Wolfe 1981). Growth on mannitol suffers from a very long lag phase unless the pre-culture was also grown on mannitol (Moon et al. 2019), and even in adapted cells growth is slower than on glucose. This long lag phase suggests mannitol is a less-favored growth substrate, and to investigate this further wild-type *T. kivui* cells were pre-cultured in complex media containing either mannitol or glucose, then passaged into complex media with a mixture of both glucose and mannitol (**Fig. 1**). In both cases cells rapidly consumed the glucose in the media, but, cells pre-cultured on glucose exhibited very strong diauxic (di-phasic) growth: upon depletion of glucose growth stopped completely, and did not resume until several hours later when mannitol consumption finally began (**Fig. 1A**). While this clearly indicates the presence of a carbon catabolite repression system in *T. kivui*, in cells pre-grown on mannitol there was still some consumption of mannitol even before the glucose was completely consumed, and cells started rapidly consuming mannitol as soon as glucose was depleted (**Fig. 1B**). The use of cells pre-cultured on mannitol may explain why previous studies (Moon et al. 2019) did not detect the catabolite repression evident here.

**Fig. 1.**
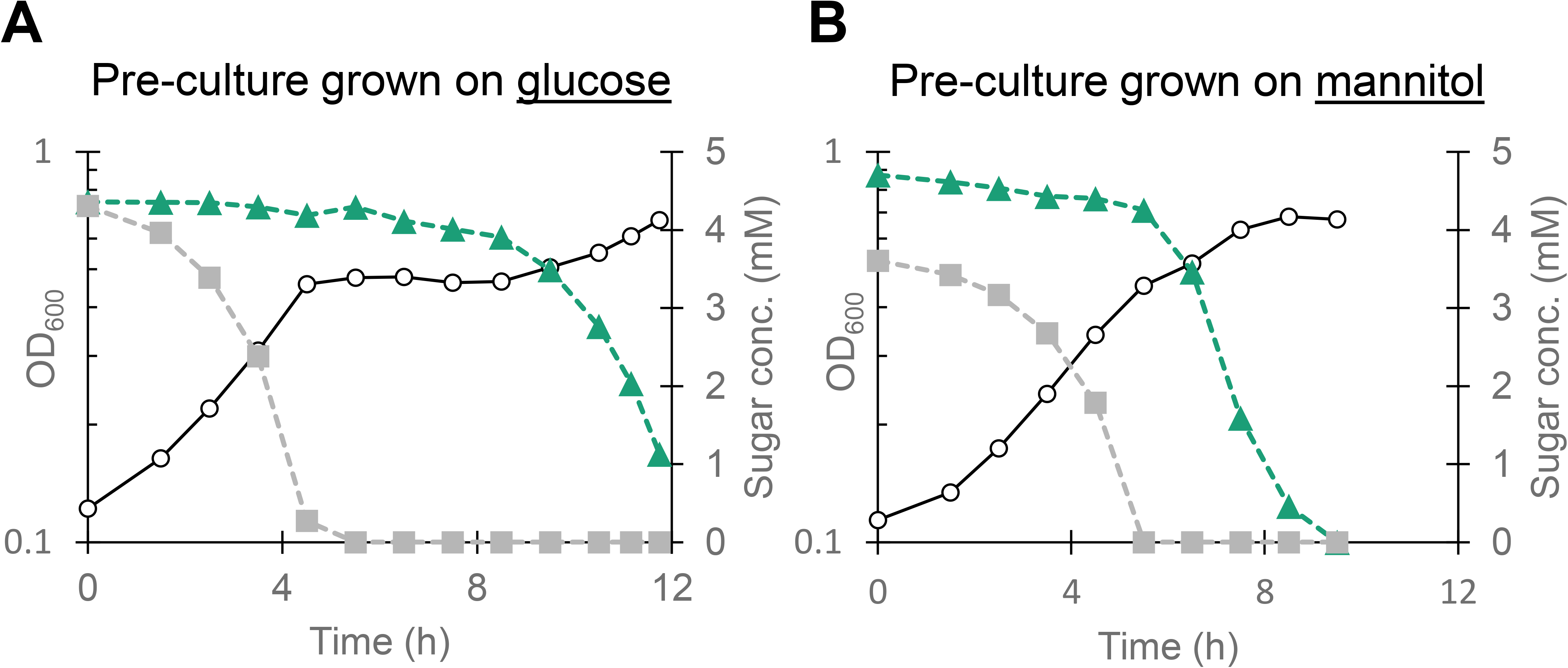
Consumption of mannitol is repressed by glucose in *T. kivui*. Cells pre-cultured on either (A) glucose, or (B) mannitol were passaged onto complex medium at 66°C containing both glucose and mannitol (5 mM each). Optical density is shown in open circles, glucose concentration in gray squares, and mannitol concentration in green triangles. One representative sample of biological duplicates is shown.

### Expression of mannitol and fructose genes on mixed sugars

Since the genes *mtlD* (Tkv_c02860) and *fruK* (Tkv_c23150) are known to be essential for growth on mannitol and fructose, respectively, expression levels of these two genes and *slp* were measured on various sugars and sugar mixtures using qPCR (**Fig. 2A**). As expected based on the RNA-sequencing data, expression of *slp* was extremely high under all growth conditions, while *mtlD* was up-regulated during growth on mannitol. Maximal expression of *mtlD* was on mannitol, at 2.5-fold higher than the reference *gyrA* gene, while minimal expression was on glucose at 8-fold below the reference gene. This implies *mtlD* is roughly 20-fold up-regulated on mannitol relative to growth on glucose, whereas RNAseq indicated genes of this operon were up-regulated between 4 and 7-fold (**Table 1**). In line with the catabolite repression observed in the sugar consumption experiments above, the addition of glucose eliminated the mannitol-induced up-regulation.

**Fig. 2.**
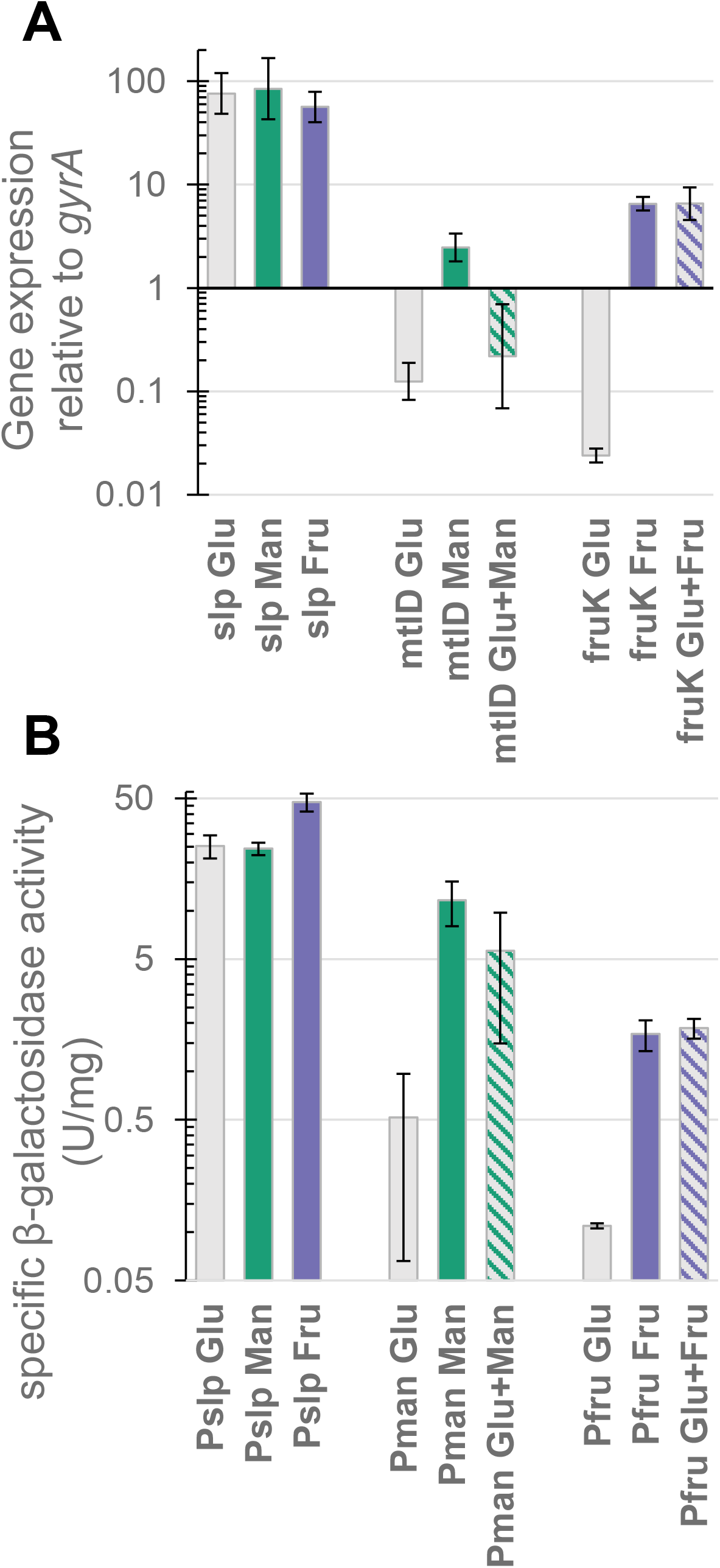
Genes of the mannitol and fructose operons are up-regulated in response to the respective sugar. (A) Expression level of *slp*, *mtlD* and *fruK* genes in wild-type *T. kivui* relative to expression of the *gyrA* gene [2^(-ΔCt)]. (B) β-galactosidase activity in cell-extracts of mutant strains of *T. kivui* expressing a thermostable β-galactosidase under the control of the three promoters being tested. Cells were grown in defined or complex medium on various sugars (30 mM) as indicated by bar color (gray = glucose, green = mannitol, purple = fructose, while hashed bars are mixtures of two sugars). Cells were grown at 66°C and β-galactosidase activity assayed at 65°C. Each bar represents average ± SD of at least two biological replicates.

The *fruK* gene exhibited even higher induced expression (6-fold above *gyrA*) in cells grown on fructose, and even lower basal expression (40-fold lower than *gyrA*), for an up-regulation on fructose more than 250-fold relative to growth on glucose (**Fig. 2A**). Up-regulation of *fruK* is also evident in cultures grown on a mixture of glucose and fructose, indicating fructose utilization is not repressed by the presence of glucose. Additional tests with all combinations of sugars used by *T. kivui* (glucose, fructose, mannitol and mannose) indicated that *mtlD* was repressed by all other sugars, while *fruK* expression was unaffected (**Fig. S2**). Since *fruK* expression combined low-leakiness, high induced expression and ability to be used in combination with other sugars, the promoter of the fructose operon seemed to be the best candidate for a sugar inducible promoter, while the mannitol promoter is of interest due to its ability to be repressed by simple addition of another sugar substrate.

To confirm that the transcriptional responses in the native genes detected by qPCR translate to heterologous protein expression, the β-galactosidase (β-gal) of the extremely thermophilic, plant biomass deconstructing *Caldicellulosiruptor bescii* (Yang et al. 2010; Basen et al. 2014) was cloned into the *T. kivui* genome as a reporter gene, under the control of the three promoters being tested (referred to as strains P_slp_, P_fru_ and P_man_ based on which promoter they contain). The promoter strains were cultured on various sugars and cells harvested for β-gal activity assays to determine the level of protein expressed (**Fig. 2B**). No β-gal activity could be detected in wild-type cells under any growth condition. Activity of the reporter gene with P_slp_ and P_man_ followed trends similar to the transcription levels of the native genes the promoters were derived from (**Fig. 2A**). Reporter activity in P_man_ cells was around 20-times higher when grown on mannitol than on glucose, while activity in P_slp_ cells was around 2.5 times higher than the highest activity in strain P_man_, and showed minimal dependence on growth sugar. However, despite qPCR results showing transcription levels of the *fruK* gene could be several times higher than *mtlD*, the highest activity obtained by P_fru_ cells was less than 20% as high as the highest activity for P_man_, although this still represents a nearly 20-fold increase compared to the very low activity on glucose. The strong repression of *mtlD* transcription by the addition of sugars other than mannitol detected with qPCR (**Fig. 2A** and **Fig. S2**) is less evident here, but average β-gal activity of cells grown on glucose + mannitol is still less than half that of cells grown on mannitol alone.

Since β-gal should allow growth on lactose, which *T. kivui* cannot natively utilize, the strains were also tested for their ability to grow on lactose. Wild-type and P_fru_ cells do not grow on lactose, while P_slp_ cells reached an optical density of 1 within a few days (**Fig. S3**). For strain P_man_, growth was dependent on the pre-culture, with cells grown on mannitol and already expressing the β-gal able to grow very slowly (reaching an OD of 0.5 after around 1 week), while cells passaged from glucose exhibited no growth. These results agree perfectly with the relative β-gal activity levels shown in **Fig. 2B**. The, roughly linear growth rates visible in **Fig. S3** likely result from a limitation in lactose hydrolysis. The β-gal lacks a signal peptide for export, and *T. kivui* is not expected to import lactose. Therefore, cell growth is only possible once enough enzyme leaks out of the cytoplasm, or enough cells lyse open, that extracellular lactate can be cleaved.

### Investigating operon arrangement

Interestingly, the gene encoding the S-layer protein *slp* is directly upstream of the fructose operon. Therefore, the native control of the genes involved in fructose metabolism may be influenced by *slp*, or a portion of the promoter region may have been missed during cloning, leading to the observed unexpectedly low β-gal activity in the strain P_fru_. To confirm the organization of the fructose and mannitol uptake operons, cDNA generated from cells grown on glucose, fructose, or mannitol was amplified with the primers depicted in **Fig. 3A**, with genomic DNA as a positive control and RNA as negative control. The lack of a PCR product with a forward primer binding in *slp* (BZq05) and a reverse primer binding in *fruR* (the first gene of the fructose operon, BZq112) clearly indicated that *slp* and *fruR* are expressed as separate transcripts, while *fruR* (BZq111) and *fruK* (BZq99) formed an operon as expected (**Fig. 3B**). The P_man_ promoter is preceded by a *levR* gene annotated as a transcriptional regulator, but here again there was no evidence that this gene was co-transcribed with the first gene of the mannitol operon, which encodes portions of enzyme II (EII) of the PTS system (primers BZ141 & BZq57). In contrast, *mtlR* and *mtlD* are clearly part of an operon (BZq59 & BZq20). Since the recombinant strains also contain the selective marker *pyrE* upstream of the reporter gene, primers were also used to test whether there was any read-through from the X514 gyrase promoter P_gyrX514_ used to express *pyrE*. There was no evidence that the selective marker and reporter gene were co-transcribed (BZq07 & BZq87) indicating that the terminator sequence included after *pyrE* was effective, although it is worth noting that this method only detected co-transcription of genes of the fructose and mannitol operons when cells were grown on the inducing sugar, meaning that lower levels of read-through from *pyrE* (equivalent to the non-induced activity of P_man_ or P_fru_) could be occurring without being detected here. To avoid any possibility of read-through from the *pyrE* gene, its direction was reversed in cloning constructs used for the knock-down constructs described in the next section.

**Fig. 3.**
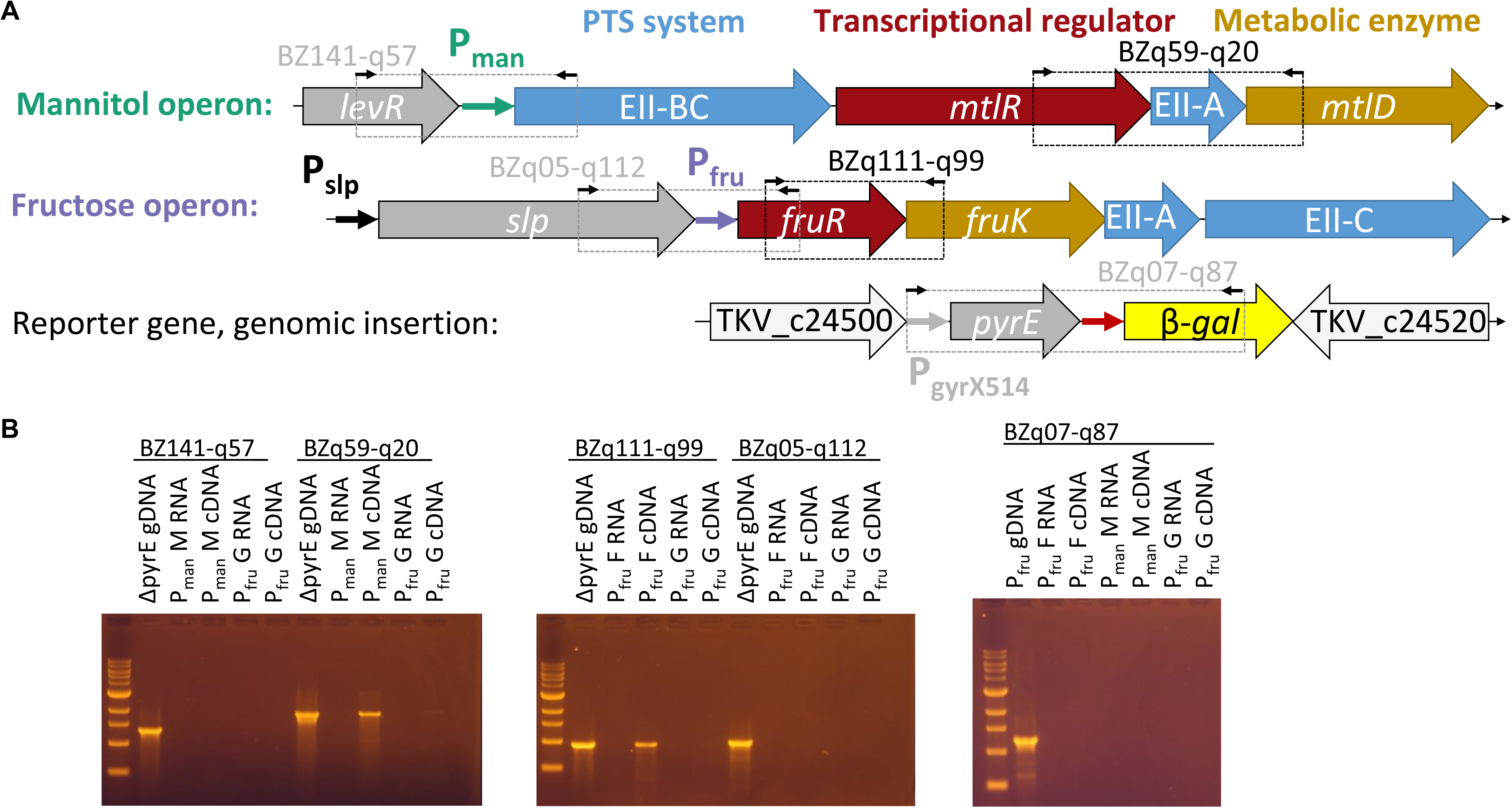
Operon arrangement and co-transcription tests. (A) Arrangement of mannitol and fructose operons in the genome of *T. kivui*, as well as the site of genomic insertion of strains with the reporter gene. Colors of large arrows indicate gene function with gene name or product inside the arrow, with red=regulator, orange=metabolic enzyme, and blue genes encoding portions of enzyme II of the PTS system. Medium arrows are promoter regions used in this work, and small black arrows indicate primer binding sites, with boxes indicating the PCR product of two primers. (B) DNA agarose gels of co-transcription tests. Each lane is labeled with the strain from which RNA or DNA was extracted, with RNA and cDNA samples also specifying the sugar cells were grown on. A band in the cDNA lane indicates co-transcription, while gDNA serves as positive control and RNA (no reverse-transcription) as negative control.

### Use of the inducible promoters to “knock-down” expression of an essential gene

Although a knockout strain of the Ech2 operon has been recently described (Baum et al. 2024), the Ech1 operon could not be knocked-out after multiple attempts, suggesting that the function of Ech1 is essential for *T. kivui*. Therefore, we decided to replace the promoter of the Ech1 operon (450 bp upstream of gene TKV_c01230) with the characterized sugar inducible promoters, generating strains P_fru_Ech1 and P_man_Ech1 (**Fig. 4A**). Cells were plated on selective medium containing the inducing sugar, and the resulting transformants characterized for their ability to grow on the inducing sugar, or glucose.

**Fig. 4.**
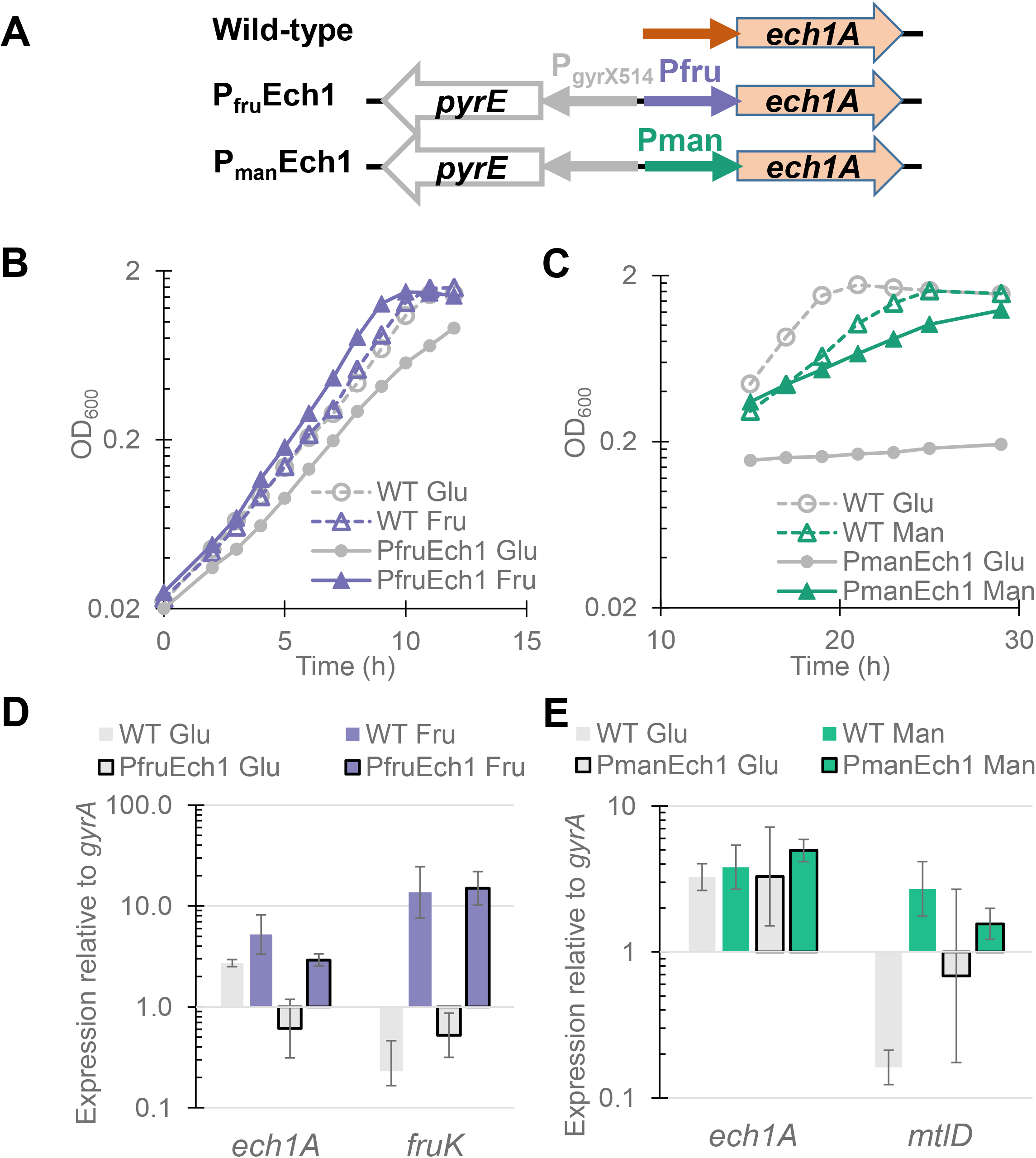
Knockdown of Ech1 operon reduces growth rate. (A) the Ech1 promoter region of *T. kivui* wild-type and the two knockdown strains, showing the reversed *pyrE* gene used for selection of mutants. Optical density of P_fru_Ech1 (B), and P_man_Ech1 (C) on defined medium at 66°C with either glucose (gray circles), fructose (purple triangles) or mannitol (green triangles), all at 30 mM. Open shapes and dashed lines indicate wild-type controls, filled shapes and solid lines are knock-down strains. A single representative growth curve of at least biological triplicates is shown. Gene expression levels relative to the *gyrA* reference gene for P_fru_Ech1 (D) and P_man_Ech1 (E), during growth on glucose (gray), fructose (purple) or mannitol (green), bars without black outline are wild-type controls. For qPCR each bar represents average and standard deviation of at least 3 biological replicates.

A growth defect was evident for both Ech1 knock-down strains when grown on the non-inducing sugar glucose (**Fig. 4B** and **C**). Wild-type *T. kivui* always grows slightly faster on fructose than glucose, but this difference was more pronounced in P_fru_Ech1 cells, which grew much worse than wild-type on glucose, but as well as, or even better than, wild-type cells on fructose (**Fig. 4B**). Growth of *T. kivui* wild-type is slower on mannitol than on glucose (Baum et al. 2024), and the P_man_Ech1 strain grew slower than wild-type on mannitol, while growth on glucose was barely detectable (**Fig. 4C**). Because of the much slower growth rates for P_man_Ech1, cells were inoculated to a lower starting OD the night before the growth experiment and allowed to grow for approximately 12 hours before monitoring began. Despite the slow growth rate, the P_man_Ech1 strain grown on glucose did eventually reach final OD_600_ values above 1.0, but this required incubation for 4 to 5 days.

Quantitative PCR was used to quantify transcription of *ech1A*, which confirmed that expression was much lower in P_fru_Ech1 during growth on glucose, and roughly as high as the wild-type during growth on fructose (**Fig. 4D**). Expression of *ech1A* measured by qPCR was similar to wild-type levels for P_man_Ech1 grown on mannitol, while expression varied wildly for P_man_Ech1 on glucose (**Fig. 4E**, note the large error bars for P_man_Ech1 Glu), which is likely a result of the very slow growth on this condition. Cellular stress responses can complicate transcriptional analysis by reducing RNA quality (Mukherjee et al. 2012), while very slow growth may have disrupted the relationship between expression levels of the reference gene (*gyrA* is tied to growth) and the gene of interest. This experiment indicated that Ech1 is important in redox metabolism during growth on sugars in *T. kivui* wild-type, since both strains exhibit a growth defect when *ech1* expression is knocked-down by growth on glucose. However, the lower levels of Ech1 still seem to be sufficient to support growth, and it is not clear why the phenotype is much more pronounced in the P_man_Ech1 strain.

The Wood-Ljungdahl pathway (WLP) appears to be essential in *T. kivui* during all types of growth, since a *T. kivui* mutant devoid of the first enzyme of the WLP, the formate-producing hydrogen-dependent carbon dioxide reductase (HDCR) was incapable of growth under all conditions unless formate was supplemented (Jain et al. 2020). Most genes of the WLP in *T. kivui* cluster together (Poehlein et al. 2015), and the RNAseq data suggests a large gene cluster starting with formyltetrahydrofolate (THF) synthetase (*fhs*, TKV_c19930), the next step in the pathway after HDCR, is co-regulated. Therefore, the promoter region (400 bp upstream) of *fhs* was replaced with the mannitol promoter to generate strain P_man_WLP.

The P_man_WLP strain grew at essentially the same rate on the inducing sugar (mannitol) and on glucose, while the wild-type control shows the expected faster growth on glucose (**Fig. 5A**). Due to the generally slower growth rates of P_man_WLP, the same overnight incubation strategy as for P_man_Ech1 was used for these growth experiments, meaning data collection started at around 14 hours. Expression of *fhs* was reduced 40-fold in the knock-down strain on glucose relative to growth on mannitol, while on mannitol *fhs* expression in the mutant was as high as in the wild-type (**Fig. 5B**). It is interesting that the P_man_ promoter was capable of expressing *fhs* at native levels, despite the fact that this is 8-fold higher than the wild-type’s expression of *mtlD* during growth on mannitol, suggesting the genomic context of *fhs* is important for its high expression.

**Fig. 5.**
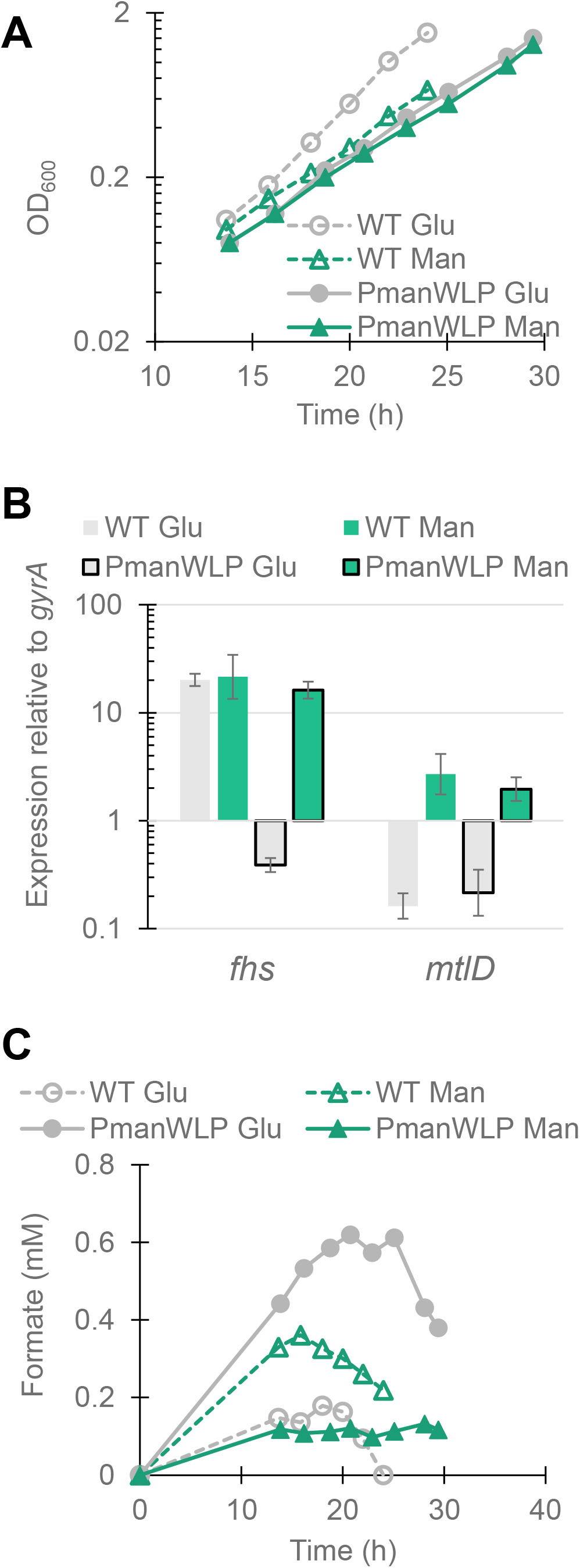
Knockdown of the WLP operon leads to formate accumulation. (A) Optical density, (B) gene expression and (C) extracellular formate concentrations of P_man_WLP growing at 66°C on defined medium with glucose (gray) or mannitol (green). A single representative growth curve of at least biological triplicates is shown, open shapes and dashed lines indicate wild-type controls, filled shapes and solid lines are the P_man_WLP strain. Quantitative PCR results are the average of at least biological triplicates, gray bars indicate growth on glucose, green on mannitol, bars with no border are wild-type, and a black border indicates P_man_WLP.

Since the genes of the WLP-operon (starting with *fhs*) consume formate generated as an intermediate by HDCR, these cells were expected to accumulate formate under the knock-down condition (growth on glucose). HPLC measurements confirmed that low concentrations of formate accumulated in the medium under all growth conditions, peaking in late exponential phase before gradually declining (**Fig. 5C**). P_man_WLP accumulated significantly more formate (p < 0.001) growing on glucose (0.61 ± 0.04 mM) compared to mannitol (0.12 ± 0.07 mM). Glucose grown P_man_WLP also produced more formate than wild-type cells. Wild-type *T. kivui* produces small quantities of formate during growth, but most reports focus on experiments with concentrated resting cells, often utilizing gases, for which formate concentrations up to 100 mM have been reported (Schwarz and Müller 2020). Although data on actively growing cells utilizing heterotrophic substrates is limited, the sub-millimolar concentrations detected here probably reflect small amounts leaking out during growth, which cells then re-uptake and utilize near the end of the growth phase. Therefore, the 0.6 mM peak detected in supernatants of P_man_WLP cells grown on glucose likely resulted from much higher intracellular concentrations.

## Discussion

Previous recombinant gene expression in *T. kivui* has always relied on one of a handful of constitutive promoters such as P_slp_, promoter of the s-layer protein (Tkv_c23170), P_gyrX514_ from the gyrase gene of the closely related *Thermoanaerobacter sp.* X514 and P_kan_, the native promoter of the *Staphylococcus aureus* kanamycin resistance gene used in plasmid pMU131. Expression levels with P_slp_ tend to be very high, while P_gyrX514_ and P_kan_ result in weaker expression that is nonetheless adequate to allow growth on selective media when controlling the appropriate marker gene (*kanR* or *pyrE*) (Basen et al. 2018; Jain et al. 2022). More recently, a broader array of promoters was characterized based on their ability to induce expression of a fluorescent reporter gene (Hocq et al. 2023). These results confirmed the strength of the P_slp_ and P_gyrX514_ promoters, but found only minimal activity of the P_kan_ promoter, despite the fact that this promoter is known to function in *T. kivui*. An additional strong constitutive promoter from the native phosphotransacetalase gene (P_pta_) was also identified, but no activity was detectable for any of the hypothetical inducible promoters tested, including P_fru_. It seems likely that P_fru_ and P_kan_ activity were simply too weak to be detected due to the limited sensitivity of the fluorescent system, particularly at higher temperatures. Therefore, the higher sensitivity of the beta-galactosidase reporter gene used here is advantageous for evaluating promoters of intermediate strength, and was even capable of quantifying the very low un-induced activity of P_man_ and P_fru_ (**Fig. 2B**).

One obvious limitation of the inducible promoters identified here is that sugar must be provided for gene induction, which will have limited utility in industrial applications converting C1-containing gasses to carbon products. Even in their induced states they are also less strong than current constitutive promoters, with induced P_man_ expression of the reporter gene approaching 50% of the activity with P_slp_. Therefore, future work will continue to search for additional promoters that can be induced in the absence of organotrophic growth substrates.

The unexpectedly low reporter gene activity of P_fru_ was disappointing given the promising qPCR data. For this study, the intergenic sequence upstream of a particular gene was selected as its “promoter region”, but this length varied depending on the gene in question, with P_fru_ at 170 bp being the shortest. Promoter function is known to be affected by the surrounding genomic context, both upstream and downstream of transcription initiation, and shorter or “minimal” promoters appear to be particularly sensitive (Davis, Rubin, and Sauer 2011). Therefore, the possibility remains that more careful selection of the promoter region could lead to a strong fructose inducible *T. kivui* promoter. Since promoter strength can also vary with the specific gene being expressed, it is possible that activity with a different reporter gene would be stronger than observed here with *C. bescii* β-galactosidase.

The relatively minor difference in growth rates between induced and non-induced P_fru_Ech1 was surprising, since qPCR indicated expression of Ech1 was 5-fold lower on glucose. This conflicts with the much more dramatic phenotype of P_man_Ech1, which was barely capable of growth on glucose, implying an important role for the Ech1 complex. Further characterization of these strains will be required to fully elucidate the role of Ech1 in the energy metabolism of *T. kivui*.

Since formate concentrations clearly indicate a bottleneck in the WLP when growing on glucose, the virtually identical growth rate of P_man_WLP on glucose relative to mannitol is also puzzling (**Fig. 5**). This, combined with the minimal difference in growth rates of P_fru_Ech1, resembles growth behaviors observed in the ΔEch2 strain grown on pyruvate (Baum et al. 2024). In that study, ΔEch2 cells initially took multiple days to grow on pyruvate, but after several passages were able to grow as well as the wild-type. This may have resulted from an adaptive mutation similar to one seen in CO-adapted cells (a truncation of the *hycB3* gene) causing suppression of the knockdown phenotype. If a suppressor mutation is also the case here, whole genome sequencing would be required to identify the site of the adaptation. It is also possible that, contrary to expectations, Ech1 and Fhs are not essential for growth on sugars, so future studies will further characterize the ability of the knockdown strains to grow autotrophically on H_2_/CO_2_.

In this study we have identified two sugar inducible promoters that function in *T. kivui*, and confirmed they were capable of regulating expression of a heterologous reporter gene as well as native metabolic enzymes. The described promoters have applications in both basic research to improve our understanding of acetogenesis at high temperatures, as well as in applied metabolic engineering efforts to utilize *T. kivui* for industrial applications.

## Materials and methods

### Strains and Growth Conditions

*Thermoanaerobacter kivui* wild-type (DSM2030) and Δ*pyrE* [TKV002 from (Basen et al. 2018)] strains, as well as *Caldicellulosiruptor bescii* (DSM 6725) cells were revived from 25% wt glycerol stocks stored at - 70°C. A full list of strains is in **Table S2.** All cell growth experiments were carried out at 66°C in water baths or incubators.

For routine growth of *T. kivui*, cells were grown in a slightly modified version of DSMZ 171 medium (Basen et al. 2018). Complex medium contained 2 g/L yeast-extract, while a defined medium for selection of mutants based on the Δ*pyrE* parent strain lacked yeast-extract, and a solid version of both media was made by addition of 1.5% wt bacto agar. Cells were supplemented with 30 mM of appropriate growth sugar (glucose, fructose, or mannitol) from anoxic 1 M stock solutions, unless another concentration is specified. *C. bescii* cells were cultured in DSMZ 516 medium, supplemented with 20 mM cellobiose from a 0.5 M anoxic stock. For liquid cultures, either Hungate tubes with 5 mL culture volume, or serum bottles with 25 or 50 mL culture volume were used.

To test growth on lactose, wild-type and P_slp_ cells were pre-grown on glucose, while P_fru_ and P_man_ cells were cultured either on glucose or their respective inducing sugar, before passaging into Hungate tubes containing 30 mM lactose, and monitoring optical density for one week.

Growth experiments of knock-down strains were performed in defined medium with 30 mM of either glucose or the inducing sugar, inoculated from cells grown under the same conditions. For the slower growing strains P_man_WLP and P_man_Ech1, cells were inoculated to a low starting OD_600_ (0.005) the night before the experiment, and incubated overnight, before monitoring growth during the following day.

### RNA-sequencing

Wild-type *T. kivui* cells were grown in triplicate in complex medium containing either 25 mM glucose or 25 mM mannitol. One milliliter of cells was harvested in mid- to late-exponential phase (OD= 1.1 for glucose, 1.3 for mannitol), rapidly cooled, and kept frozen until RNA extraction.

Harvested cells were re-suspended in 800μl RLT buffer (RNeasy Mini Kit, Qiagen) with β-mercaptoethanol (10μl ml^-1^) and cell lysis was performed using a laboratory ball mill. Subsequently, 400μl RLT buffer (RNeasy Mini Kit Qiagen) with β-mercaptoethanol (10μl ml^-1^) and 1200μl 96% (vol./vol.) ethanol were added. For RNA isolation, the RNeasy Mini Kit (Qiagen) was used as recommended by the manufacturer, but instead of RW1 buffer RWT buffer (Qiagen) was used in order to isolate RNAs smaller than 200 nucleotides also. To determine the RNA integrity number the isolated RNA was run on an Agilent Bioanalyzer 2100 using an Agilent RNA 6000 Nano Kit as recommended by the manufacturer (Agilent Technologies, Waldbronn, Germany). Remaining genomic DNA was removed by digesting with TURBO DNase (Invitrogen, Thermo Fischer Scientific, Paisley, UK). The Illumina Ribo-Zero rRNA Depletion Kit ((Illumina Inc., San Diego, CA, USA) was used to reduce the amount of rRNA-derived sequences.

For sequencing, the strand-specific cDNA libraries were constructed with a NEB Next Ultra II Directional RNA library preparation kit for Illumina and the NEB Next Multiplex Oligos for Illumina (96) (New England BioLabs, Frankfurt am Main, Germany). To assess quality and size of the libraries samples were run on an Agilent Bioanalyzer 2100 using an Agilent High Sensitivity DNA Kit as recommended by the manufacturer (Agilent Technologies). Concentration of the libraries was determined using the Qubit® dsDNA HS Assay Kit as recommended by the manufacturer (Life Technologies GmbH, Darmstadt, Germany). Sequencing was performed on the HiSeq4000 instrument (Illumina Inc., San Diego, CA, USA) using HiSeq4000 Reagent Kit sequencing in 50bp Single Read mode. For quality filtering and removing of remaining adaptor sequences, Trimmomatic-0.39 (Bolger, Lohse, and Usadel 2014) and a cutoff phred-33 score of 15 were used.

The mapping against the reference genomes of *Thermoanaerobacter kivui* DSM ^T^ 2030 was performed with Salmon (v 1.5.2) (Patro et al. 2017). As mapping back-bone a file that contains all annotated transcripts excluding rRNA genes and the whole genome of the reference as decoy was prepared with a k-mer size of 11. Decoy-aware mapping was done in selective-alignment mode with ‘–mimicBT2’, ‘– disableChainingHeuristic’and‘–recoverOrphans’flags as well as sequence and position bias correction and 10 000 bootstraps. For–fldMean and–fldSD, values of 325 and 25 were used respectively. The quant.sf files produced by Salmon were subsequently loaded into R (v 4.3.2) (R Core Team 2020) using the tximport package (v 1.28.0) (Soneson, Love, and Robinson 2016). DeSeq2 (v 1.40.1) (Love, Huber, and Anders 2014) was used for normalization of the reads and fold change shrinkages were also calculated with DeSeq2 and the apeglm pack-age (v 1.22.1) (Zhu, Ibrahim, and Love 2019). Genes with a log2-foldchange of +2/-2 and a p-adjust value ≤ 0.001 were considered differentially expressed. Raw reads have been deposited in the Sequence Read Archive as SRR28537724-SRR28537729.

### Generation of *T. kivui* mutant strains

All mutants were based on the Δ*pyrE* parent strain TKV_MB002, and used methods based on those previously described (Basen et al. 2018). For reporter gene strains, linear PCR cloning constructs contained flanking regions of a previously identified quiescent region of the genome (between genes Tkv_c24500 and Tkv_c24520) with a copy of the native *T. kivui pyrE* gene under the control of the *T. sp.* X514 gyrase promoter, followed by a terminator sequence and the reporter gene (Athe_1927) under control of one of the tested promoters. The resulting numbered mutant strains are referred to in the text by the promoter used to control the reporter gene (strain TKV_MB141 = P_slp_, TKV_MB145 = P_man_, and TKV_MB142 = P_fru_). All mutant strains retain *pyrE* and are therefore capable of growth on defined media. Cloning fragments were amplified with Q5 high-fidelity polymerase (New England Biolabs, Ipswich, MA, USA), while OneTaq polymerase (New England Biolabs) was used for screening of plasmids and mutant strains. A full list of strains is in **Table S2**, sequences of all primers are listed in **Table S3**.

For generation of strain P_slp_, the reporter gene was amplified from *C. bescii* genomic DNA with primers BZ170 and BZ171, while a plasmid backbone containing the flanking regions, *pyrE* gene, P_slp_ (the 220 bp upstream of *slp*), and an N-terminal His6-tag, was amplified from pJM009 (Nissen et al. 2024) with primers BZ172 and BZ173. The resulting PCR fragments were combined to generate plasmid pTkv141 using the NEBuilder HiFi DNA Assembly Kit (New England Biolabs), and transformed into chemically competent DH5 *E. coli* cells.

For generation of strain P_fru_, the native promoter of the *T. kivui* fructose uptake operon, consisting of 170 bp upstream and the sequence coding for the first six amino acids of Tkv_c23160, was amplified with primers BZ176 and BZ177, while the entirety of pTkv141, except for the P_slp_ promoter and N-His-tag, was amplified with primers BZ174 and BZ175. The PCR fragments were combined as described above to generate pTkv142.

For generation of strain P_man_, the native promoter of the *T. kivui* mannitol operon, consisting of 250 bp upstream and the sequence of the first four amino acids of Tkv_c02830, was amplified with BZ184 and BZ185, and the pTkv141 backbone was amplified with BZ182 and BZ183. The PCR fragments were combined to generate pTkv145.

Linear cloning fragments were amplified using primers BZ168 and BZ169, which bind at the outer edges of the flanking regions used in all three reporter gene plasmids. For transformation 1 µg of PCR product was added to Hungate tubes containing 5 mL of complex medium, inoculated with Δ*pyrE* cells, and allowed to grow overnight. A large inoculum from this culture (1 mL) was then passaged into a fresh Hungate tube containing defined medium, and allowed to incubate for several days until growth was evident. The cells which grew in defined liquid medium were plated onto defined solid medium and allowed to grow for at least 5 days. Resulting colonies were picked again into defined medium and, once grown, screened for transformants by PCR, using 2 uL culture as template, and primers LH28 and LH29. Some colonies screened mixed at this stage, with both parent strain (Δ*pyrE*) and transformant bands evident on DNA gels, so a second or even third round of plating was required to obtain contaminant-free mutants. To confirm the identity of mutant strains, and ensure no mutations were present in the promoter region, PCR products were sent to LGC Genomics (Berlin, Germany) for sequencing.

The method to generate strains with the native Ech1 and Wood-Ljungdahl operon promoters substituted for the inducible P_man_ or P_fru_ was similar to the reporter gene strains, but with flanking regions to knock out the native promoter of the gene being targeted (*ech1A* or *fhs*), and the direction of the P_gyrX514_-*pyrE* cassette was reversed, so that it pointed away from the inducible promoter. The strains are numbered based on the plasmid used to generate them, but in the text are referred to by the identity of the inducible promoter and the operon it controls (TKV_MB144 = P_fru_Ech1, TKV_MB148 = P_man_Ech1, TKV_MB156 = P_man_WLP). Unless otherwise stated, plasmids were assembled using the NEBuilder-HiFi-DNA assembly kit. Screening and sequencing of transformants was as described for the reporter gene strains, but with different screening primers.

To generate plasmid pTkv144, first an intermediate plasmid pTkv134 was generated containing flanking regions to the Ech1 operon promoter. The DFR was amplified from *T. kivui* gDNA using BZ146 and BZ147, the UFR using BZ148-149, and the backbone of pMBTkv022 (Moon et al. 2019) using LH001 and LH006. Then intermediate plasmid pTkv135 was generated by amplifying the p134 backbone with flanking regions using BZ152-153, and P_fru_ amplified for insertion between the flanking regions with BZ150-151. Then intermediate plasmid p143 was generated from the pJM009 backbone and pTkv135 insert by digesting both plasmids with BamHI/XbaI, gel extracting the appropriate-size bands, and ligating. Finally, pTkv144 was generated by amplifying the p143 backbone including flanking regions and P_fru_ with BZ178-179, and combining with the backwards P_gyrX514_-*pyrE* cassette from pJM009 using BZ180-181. To generate p148, the entirety of pTkv144 except for the P_fru_ promoter was amplified using BZ196-197. The P_man_ promoter for insertion into this backbone was amplified from *T. kivui* gDNA using BZ198-199. Cloning constructs from p144 and p148 were amplified using BZ190-191, and the resulting linear PCR products used to transform *T. kivui* Δ*pyrE* cells. Primers BZ154 and BZ155 were used to screen for successful transformation.

To generate pTkv156, first an intermediate plasmid was generated with the whole region upstream of *fhs*, including the native promoter, DFR, and UFR, amplified from *T. kivui* gDNA using BZ215 and BZ216, and cloned into the backbone of pTkv144 amplified with LH001 and LH006. The backbone plus flanking regions of the intermediate plasmid was amplified using BZ218 and BZ239, and combined with the P_gyrX514_-*pyrE* cassette and P_man_ promoter amplified from pTkv148 using BZ219 and BZ240. The linear PCR fragment used for cloning in *T. kivui* was amplified from pTkv156 using BZ221 and BZ222. Primers BZ223 and BZq28 were used to screen for successful transformation.

### Quantitative RT-PCR

Cells (2 mL) were harvested in mid-exponential phase (OD 0.4 – 0.7), rapidly cooled on ice, centrifuged at 15,000 g for 5 minutes, and the pellets stored at -70°C until RNA extraction. RNA was extracted using the QIAGEN (Hilden, Germany) RNeasy kit with the optional on-column DNase-digestion. According to the modifications for bacterial cells recommended by the manufacturer, cells were re-suspended in lysis buffer RLT containing β-mercaptoethanol, added to tubes containing 50 mg acid-washed beads, and processed in a FastPrep-24 bead beater at 6 m/s for four 40 second cycles (MP Biomedicals, Irvine, CA, USA), before continuing with the rest of the standard protocol. The on-column DNase-digestion was extended to 30 minutes to further reduce gDNA contamination.

For synthesis of cDNA, 10 µL reactions of the Biozym cDNA Synthesis Kit (Biozym Scientific, Hessisch Oldendorf, Germany) containing 100 ng of RNA were incubated at 42°C for 30 minutes. The noRT controls consisted of RNA diluted to the same concentration in RNase-free water.

Quantitative PCR was performed with a qTower^3^ G cycler (Analytik Jena, Jena, Germany). The Biozym Blue S’Green qPCR Mix was used, with 2 µL of 10x diluted cDNA or noRT template in 10 µL reactions. The *T. kivui* gyrase subunit A (*gyrA*, Tkv_c00100) was used as reference gene, and a noRT control with the *gyrA* gene was included for each RNA extraction. Sequences of all primers used for qPCR are included in **Table S3**.

### Co-transcription tests

To determine operon structure, RNA extracted for qPCR was further treated with the TURBO DNA-*free* kit (ThermoFisher Scientific, Waltham, MA, USA) to remove traces of residual gDNA. This RNA was then reverse-transcribed, and the resulting cDNA used as template for PCR with primers binding in the upstream and downstream genes. Positive controls consisted of the same PCR reaction with gDNA as template, and negative controls with RNA as template. Primer binding sites are shown in **Fig. 5A**, and primer sequences listed in **Table S3**.

### Enzyme activity assays

Cells were grown on various sugars and sugar mixtures to a final OD of 0.8 to 1.2, and cell pellets were harvested from approximately 20 mL of cell culture and frozen until analysis. Cell pellets were re-suspended in 650 µL of lysis buffer (50 mM Tris-HCl, 100 mM NaCl, pH 7.8) and processed in a FastPrep-24 bead beater at 6 m/s for four 40 second cycles. Lysed cells were then centrifuged at 15,000 g for 5 minutes, and the supernatant (cell-extract) was transferred to a fresh tube.

Protein concentration of the cell-extract was determined by the Bradford method using the ROTI Nanoquant reagent (Carl Roth, Karlsruhe, Germany) in a 96-well plate, with BSA as standard curve.

Activity assays were carried out in 1.7 mL tubes containing 600 µL of substrate solution (4.67 mM pNP-β-gal, dissolved in 58 mM sodium acetate buffer, pH 5.8). Tubes containing substrate were pre-heated in a 65°C heat block for 5 minutes, then 100 µL of diluted cell-extract was added, and 100 µL samples were taken at various timepoints into wells of a 96-well plate containing 100 µL of 1 M Na_2_CO_3_ to stop the reaction. Concentration of pNP was determined using a standard curve consisting of serial dilutions of a reaction that was allowed to go to completion, and change in pNP concentration over time was used to calculate specific activity of each cell extract (µmol/min/mg).

### Monitoring growth and metabolite concentrations

Cells were cultured in serum bottles with 1 mL samples taken at various timepoints. Half of each sample (500 µL) was mixed with an equal volume of distilled water for immediate determination of OD_600_. The remaining 500 µL was acidified with 10 µL 50% sulfuric acid and stored frozen. Samples were thawed, centrifuged to remove cells, and analyzed by HPLC using an organic-acid-resin 300×8 mM column (CS-Chromatography Service GmbH, Langerwehe, Germany) at 30°C on a Shimadzu HPLC system (Kyoto, Japan) consisting of LC-20AD pump, SIL-20AC autosampler, CTO-20AC column oven, and RID-10A RI detector, with mobile phase consisting of 5 mM sulfuric acid at 0.6 mL/min. Concentrations in 10 µL injections were determined by comparison to standard curves of glucose, mannitol, fructose, and sodium formate.

## Acknowledgements and funding

Funding was provided by Deutsche Forschungsgemeinschaft (DFG, project number 394854436), and Bundesministerium für Bildung und Forschung (BMBF) project ThermoSynCon (grant numbers 031B0857A–C).

## Author Contributions

Conceptualization BZ and MB; data curation AP; funding acquisition MB, RD, and VM; investigation BZ, SM, CB, AS, and AP; supervision MB, BZ, RD, and VM; writing – original draft BZ; writing – review and editing all authors.

