## Supplemental Tables and Figures for "Knock-down of genes essential for homoacetogenic growth using sugar inducible promoters in the thermophile *Thermoanaerobacter kivui*"

**Table S2:** Strains used or generated in this study

| Strain | Genetic modifications | Parent strain | Reference |
|---|---|---|---|
| *T. kivui* DSM2030 | NA | NA | (Leigh, Mayer, and Wolfe 1981) |
| *C. bescii* DSM 6725$^T$ | NA | NA | (Yang et al. 2010) |
| TKV_MB002 (Δ*pyrE*) | Δ*pyrE* (TKV_c14380) | *T. kivui* DSM2030 | (Basen et al. 2018) |
| TKV_MB141 ($P_{slp}$) | →$P_{gyrX514}$*pyrE* →$P_{slp}$β-*gal* (Athe_1927)* | TKV_MB002 | This study |
| TKV_MB142 ($P_{fru}$) | →$P_{gyrX514}$*pyrE* →$P_{fru}$ β-*gal* (Athe_1927)* | TKV_MB002 | This study |
| TKV_MB145 ($P_{man}$) | →$P_{gyrX514}$*pyrE* →$P_{man}$β-*gal* (Athe_1927)* | TKV_MB002 | This study |
| TKV_MB144 ($P_{fru}$Ech1) | $P_{gyrX514}$*pyrE* ⟵ →$P_{fru}$*ech1A* (TKV_c01230) | TKV_MB002 | This study |
| TKV_MB148 ($P_{man}$Ech1) | $P_{gyrX514}$*pyrE* ⟵ →$P_{man}$*ech1A* (TKV_c01230) | TKV_MB002 | This study |
| TKV_MB156 ($P_{man}$WLP) | $P_{gyrX514}$*pyrE* ⟵ →$P_{man}$*fhs* (TKV_c19930) | TKV_MB002 | This study |

* inserted in region between TKV_c24500 and TKV_c24520

**Table S3**: Primers used in this study

| Primer | Binds | Product | **Sequence** (lower case letters = non-binding tail for plasmid assembly) |
|---|---|---|---|
| **Cloning** | | | |
| BZ146 | *T. kivui* gDNA | Ech1 promoter UFR | ctcggtacccggggatccGACAGAAGAAGAAATATACAGGGTTTCTG |
| BZ147 | | | ggtataaaggatgtcatCACACTCTCCTCGTACAGCTT |
| BZ148 | *T. kivui* gDNA | Ech1 promoter DFR | cgaggagagtgtgATGACATCCTTTATACCCATAATGTTAG |
| BZ149 | | | gcatgcctgcaggtcgactctagaCTATCAGCCAGAGCATCAAAGTAG |
| BZ150 | *T. kivui* gDNA | $P_{fru}$ | gagtgtgGGACTCTTTTATTTGCTTTAAAACAGGC |
| BZ151 | | | gggtataaaggatgtcatATCAGATATACTCATATTCATTCACATCC |
| BZ152 | pTkv134 | bb, UFR, and DFR | cctgttttaaagcaaataaaagagtccCACACTCTCCTCGTACAGCTT |
| BZ153 | | | ggatgtgaatgaatatgagtatatctgatATGACATCCTTTATACCCATAATGTTAG |
| BZ154 | gDNA of *T. kivui* (screening) | *ech1A* promoter region | GTAGAAGCAGTGGATAAAGTCATACTG |
| BZ155 | | | GACCTATAATAGAGGCGCTTAAATAGATG |
| BZ168 | Reporter gene plasmids | linear cloning fragment | TCTCTTATCACCACCTCCACA |
| BZ169 | | | GCAATTATGGCAATGCGTGG |
| BZ170 | *C. bescii* gDNA | Amplify reporter gene Athe_1927 | catcatcaccacATGGGCAAAATAAAGCTGAAAAAATTTTTGC |
| BZ171 | | | ggactaagcccccatttTTAATCTTTTAATCTTCTCAGTATCATAACCTCC |
| BZ172 | pJM009 | bb, UFR and DFR | gattaaaagattaaAAATGGGGGCTTAGTCCC |
| BZ173 | | | gctttattttgcccatGTGGTGATGATGGTGATGCATACAG |
| BZ174 | pTkv141 | bb + UFR, *pyrE*, and DFR | gaatatgagtatatctgatATGGGCAAAATAAAGCTGAAAAAATTTTTGC |
| BZ175 | | | gaatattattgtactatttcCATTTCTTTGAGATAATAAAATAAGCCCACCTG |
| BZ176 | *T. kivui* gDNA | $P_{fru}$ | cttattttattatctcaaagaaatgGAAATAGTACAATAATATTCAACTATAAAGAGTCC |
| BZ177 | | | gctttattttgcccatATCAGATATACTCATATTCATTCACATCCA |
| BZ178 | pTkv143 | bb, UFR, DFR, $P_{fru}$ | gagtaaaaataggGGACTCTTTTATTTGCTTTAAAACAGGC |
| BZ179 | | | caggaaattttaaCACACTCTCCTCGTACAGCTTG |
| BZ180 | pJM009 | *pyrE* | gaggagagtgtgTTAAAATTTCCTGCTTCCCG |
| BZ181 | | | ataaaagagtcccCCTATTTTTACTCTTCTTTCTGG |
| BZ182 | pTkv141 | bb, UFR, *pyrE*, and DFR | gtaaaacttatgcaaaatactATGGGCAAAATAAAGCTGAAAAAATTTTTGC |
| BZ183 | | | cttatcggcacctgtatCATTTCTTTGAGATAATAAAATAAGCCCACCTG |
| BZ184 | *T. kivui* gDNA | $P_{man}$ | cttattttattatctcaaagaaatgATACAGGTGCCGATAAGGATAGAATAG |
| BZ185 | | | gctttattttgcccatAGTATTTTGCATAAGTTTTACCTCCTTTTTG |

| | | | |
|---|---|---|---|
| BZ190 | Ech1 promoter plasmids | linear cloning fragment | CAGGGTTTCTGTACACGAAGC |
| BZ191 | | | CTCTTTCGTGATATCCATTGCTGC |
| BZ196 | pTkv144 | bb, UFR, pyrE, DFR | ggtaaaacttatgcaaaatactATGACATCCTTTATACCCATAATGTTAG |
| BZ197 | | | ccttatcggcacctgtatCCTATTTTTACTCTTCTTTCTGGTTAAAGC |
| BZ198 | *T. kivui* gDNA | $P_{man}$ | gaagagtaaaaataggATA CAGGTGCCGATAAGGATAGAATAG |
| BZ199 | | | ggtataaaggatgtcatAGTATTTTGCATAAGTTTTACCTCCTTTTTG |
| BZ215 | *T. kivui* gDNA | *fhs* UFR, DFR and Promoter | ctcggtacccggggatccGTTGTAACTTCTATGCCATGGGC |
| BZ216 | | | tgcctgcaggtcgactctagaGGTTATCACATAATCAGCAAGCTTC |
| BZ218 | intermediate to pTkv156 | bb, UFR and DFR | gtaaagccgggaagcaggaaattttaaGGTGGGCACACTACAAAAAACG |
| BZ239 | | | gcaaaaaggaggtaaaacttatgcaaaatactATGGCATTTAAGAGCGATATTGAGA |
| BZ219 | pTkv148 | *pyrE* and $P_{man}$ | TTAAAATTTCCTGCTTCCCGG |
| BZ240 | | | AGTATTTTGCATAAGTTTTACCTCCTTTTTG |
| BZ221 | pTkv156 | linear cloning fragment | GGCATTTGTAGTTAAGATTGCTAATGC |
| BZ222 | | | GGTTGCTATCACGCTGTTACAAC |
| BZ223 | gDNA of *T. kivui* (screening) | *fhs* promoter region | CAGGTGATGTTGTAACTTCTATGCC |
| BZq28 | | | CTCTTTTGCAAGCTCAATTCCAC |
| LH001 | pUC19 derived plasmids | bb | GGATCCCCGGGTACCGAG |
| LH006 | | | TCTAGAGTCGACCTGCAG |
| LH28 | gDNA of *T. kivui* (screening) | Insert region Tkv_c24500-520 | CAGGCTGTGATAATTTGAGAA |
| LH29 | | | GGTCACGATTTAAAGGACTTA |
| **RT-qPCR** | | | |
| **Primer** | **Gene name** | **Gene Orf** | **Sequence** |
| gyr-Fwd | *gyrA* | TKV_c00100 | CCAGTTGTGCTTCCTTCTCGATTTCC |
| gyr-Rev | | | GCGACAATGCCATCTATGACTTCTCC |
| BZq05 | *slp* | TKV_c23170 | GACATACAGAGGGCAAATGATCAAC |
| BZq06 | | | GTTAAGAAGCACTGTGTTGTCTGTG |
| BZq19 | *mtlD* | TKV_c02860 | CTGTGGACAGGATAGTACCGAATGTAG |
| BZq20 | | | CAAGGTCAGCGACAAGTTCCAC |
| BZq21 | *fruK* | TKV_c23150 | GACATAATTAGAGAAATAAAAAGCGGCG |
| BZq22 | | | CTACTGCATCTTCTAACTCCTGC |
| BZq28 | *fhs* | TKV_c19930 | CATTAAATAGATTCCCGACAGACACAG |
| BZq29 | | | CTCTTTTGCAAGCTCAATTCCAC |
| Ech1A f | *ech1A* | TKV_c01230 | CCTCTTTGCCGGTGTAATGAGTAAGG |
| Ech1A r | | | AAGCATGGTAAACGCACCCAAC |
| **Co-transcription tests** (for exact binding locations see Fig. 5) | | | |
| **Primer** | **Gene name** | **Gene Orf** | **Sequence** |
| BZ141 | *mtlA* | TKV_c02830 | CTTTTGGTGCCATAGCTATCTCTG |
| BZq07 | *pyrE* | (before βgal) | TGACATCAGGAAAACACAGCG |
| BZq57 | *levR* | TKV_c02820 | TACACTTTTGCTTGGGGAGACG |
| BZq59 | *mtlR* | TKV_c02840 | GCGGTAAATAGAGCTTGCAAGT |
| BZq99 | *fruK* | TKV_c23150 | GCATTAGGTCCTATAAAGCCGAGA |
| BZq111 | *fruR* | TKV_c23160 | GGAAAGGCGCTTAAAGATTGCA |
| BZq112 | *fruR* | TKV_c23160 | AACATTCAATATCGCACCACCG |

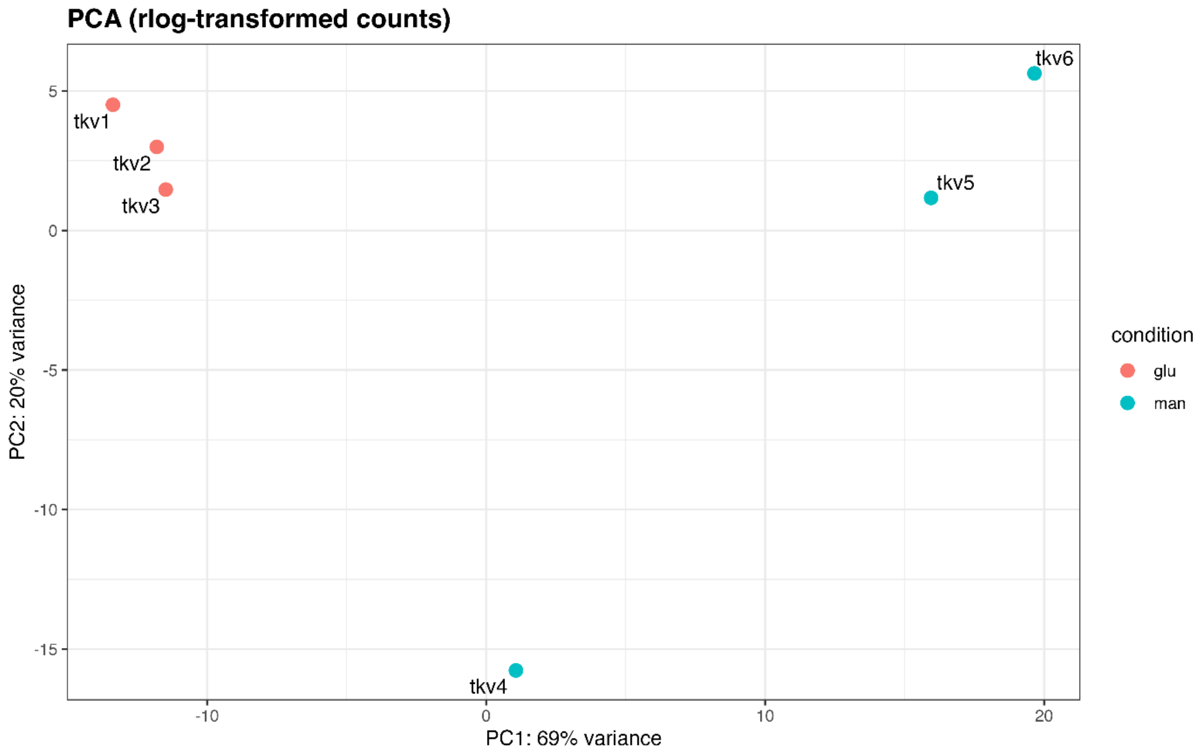


**Figure S1**: Principle component analysis of RNAseq samples. The three glucose samples (tkv1-3) are shown in red and mannitol samples (tkv4-6) in blue.

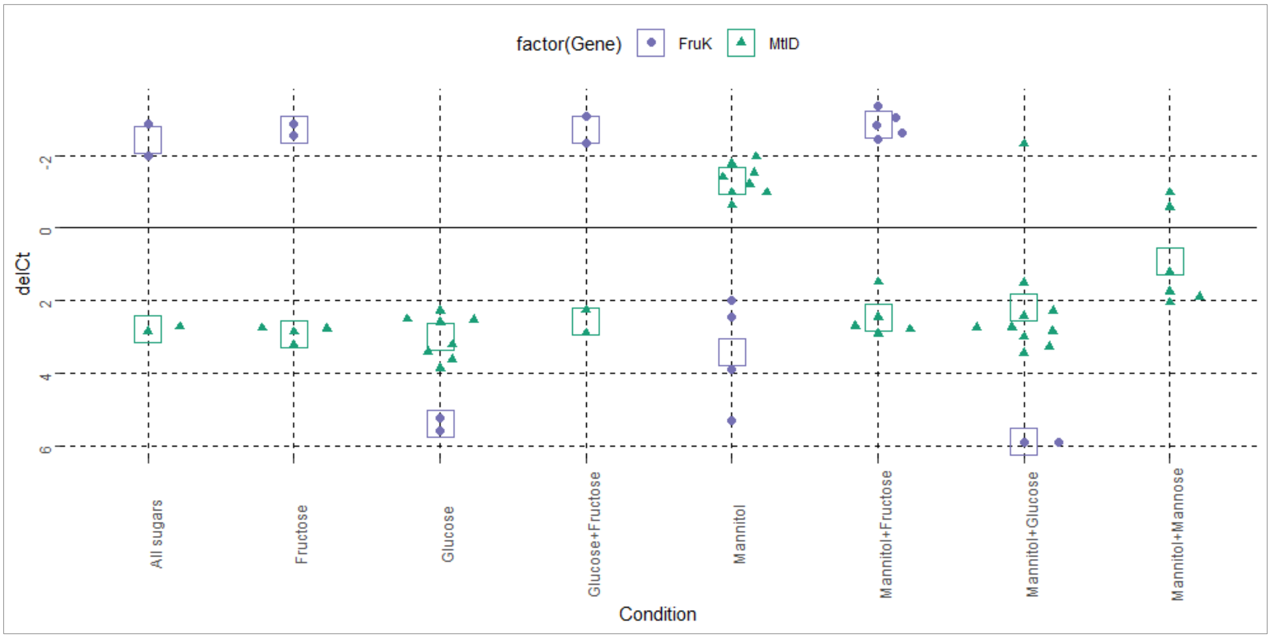


**Figure S2:** Expression levels relative to *gyrA* reference gene (ΔCt), so negative numbers indicate up-regulation (gene reached Ct earlier than the reference gene). Each *n**Ct difference = 2^*n* fold-change, as DNA concentration doubles every 1 PCR cycle. Each individual data point (*fruK* = green triangles, *mtlD* = purple circles) is a single biological replicate, open boxes show the average for each condition. Technical replicates were not performed, since previous experience found minimal variance compared to the differences between biological samples.

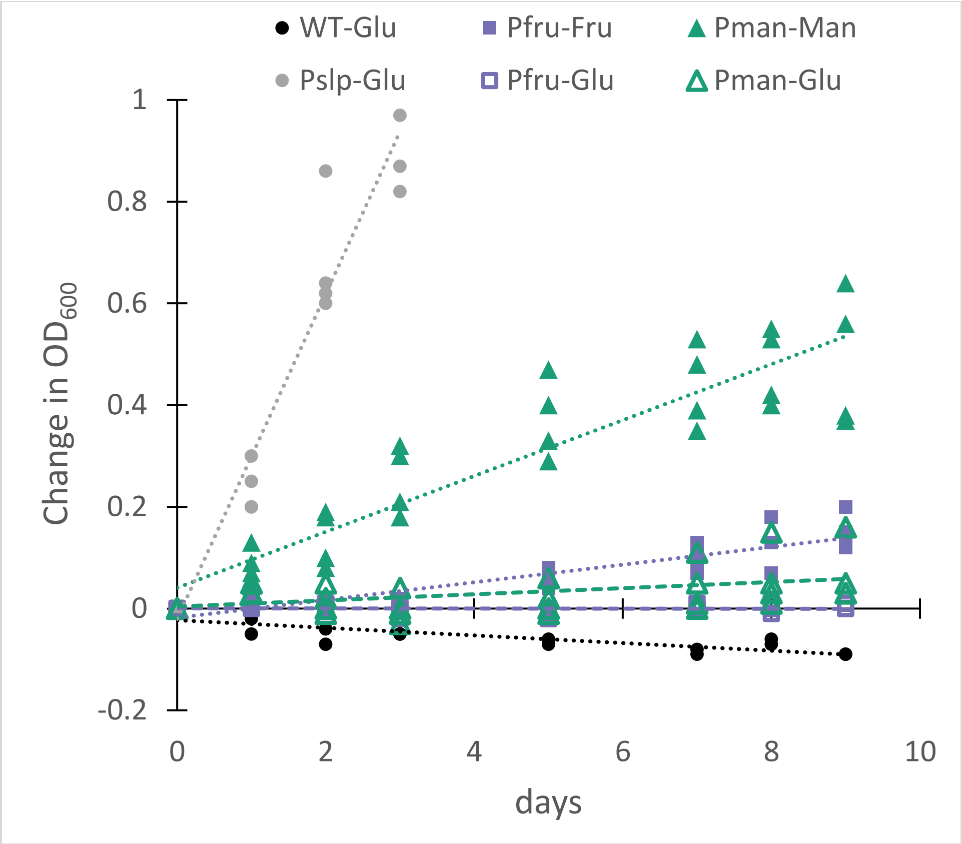


**Figure S3**: Growth of wild-type and β-galactosidase expressing strains on lactose in defined medium. For $P_{fru}$ and $P_{man}$, open shapes indicate pre-culture was grown on glucose, while filled shapes were pre-cultured on the respective inducing sugar ($P_{slp}$ was pre-cultured only on glucose). Wild-type = black circles, $P_{slp}$ = gray circles, $P_{fru}$ = purple squares, $P_{man}$ = green triangles.
